# Genome-wide CRISPR screens in T helper cells reveal pervasive cross-talk between activation and differentiation

**DOI:** 10.1101/196022

**Authors:** Johan Henriksson, Xi Chen, Tomás Gomes, Ubaid Ullah, Kerstin B Meyer, Ricardo Miragaia, Graham Duddy, Jhuma Pramanik, Kosuke Yusa, Riitta Lahesmaa, Sarah A Teichmann

**Author notes:** Equal contribution (Johan Henriksson, Xi Chen). Equal contribution (Kosuke Yusa, Riitta Lahesmaa). Johan Henriksson, Xi Chen, Tomás Gomes, Kerstin Meyer, Ricardo Miragaia, Ubaid Ullah, Graham Duddy, Jhuma Pramanik, Kosuke Yusa, Riitta Lahesmaa, Sara A Teichmann.

## Abstract

T helper type 2 (Th2) cells are important regulators of mammalian adaptive immunity and have relevance for infection, auto-immunity and tumour immunology. Using a newly developed, genome-wide retroviral CRISPR knock-out (KO) library, combined with RNA-seq, ATAC-seq and ChIP-seq, we have dissected the regulatory circuitry governing activation (including proliferation) and differentiation of these cells. Our experiments distinguish cell activation *versus* differentiation in a quantitative framework. We demonstrate that these two processes are tightly coupled and are jointly controlled by many transcription factors, metabolic genes and cytokine/receptor pairs. There is only a small number of genes regulating differentiation without any role in activation. Our study provides an atlas for the T helper cell regulatory network, pinpointing key players of Th2 differentiation and demonstrating remarkable plasticity between the diverse T helper cell fates. We provide an online resource for interactive data querying at: http://data.teichlab.org.

## Introduction

CD4+ T helper (Th) cells are a central part of the adaptive immune system and play a key role in infections, autoimmunity and tumour repression. During the immune response, Th cells become activated and differentiate from a naive state into different effector subtypes, including T helper type 1 cells (Th1), Th2, Th17 and regulatory T cells (Treg). Different subtypes have distinct functions and molecular characteristics^1^. Th2 cells are primarily responsible for eliminating helminths and other parasites and are strongly associated with allergies. Thus, a better understanding of Th2 cell development is key to combating a range of clinical conditions^2^.

Th2 differentiation is characterized by the production of the cytokines *Il4, Il5* and *Il13. In vitro, Il4* is crucial for the activation of the signalling transducer *Stat6*^3–5^, which in turn induces the Th2 master regulator *Gata3*^6–9^. As Th2 cells upregulate their key inducer, a positive feedback loop is formed. Th1 cells possess an equivalent mechanism for their defining transcription factor (TF), *Tbx21*, which represses *Gata3. Gata3* is able to inhibit *Ifng*, the main cytokine driving Th1 differentiation. Thus, the balance of the two TFs *Tbx21* and *Gata3* defines the Th1-Th2 axis^10^. There are however many genes affecting this balance, and alternative Th fates are frequently affected by overlapping sets of regulatory genes. Despite the importance of different T helper subtypes, so far only the Th17 subtype has been examined systematically^11,12^.

A major challenge in performing genetic studies in primary mouse T cells is the lack of efficient genetic perturbation tools. To date, only a small RNA interference screen has been performed *in vivo* on mouse T cells^13^. However, recently-developed CRISPR technology has the advantages of higher specificity and greater flexibility, allowing knock-out^14,15^, repression^16–18^ and activation^19–21^. Currently all existing CRISPR libraries are lentiviral-based^22,23^ and therefore unable to infect murine Th cells^24^. To overcome this limitation, we created a genome-wide retroviral CRISPR sgRNA library. By using this library on T cells from mice constitutively expressing *Cas9*, we obtained high knock-out efficiency. In addition, we established an arrayed CRISPR screening protocol that is scalable and cost-efficient.

After library transduction, we screened for and characterized genes strongly affecting Th2 differentiation. We chose *Il4, Il13, Gata3, Irf4* and *Xbp1* as our primary screen read-outs. *Il4, Il13, Gata3* are at the core of the Th2 network^10^ while *Irf4* and *Xbp1* have been suggested to have supporting roles in keeping the chromatin accessible and in overcoming the stress response associated with rapid protein synthesis during T cell activation^25–27^. Selected genes discovered by the screen were validated in individual knock-outs and assayed by RNA-seq. To place the discovered genes into the context of Th2 differentiation, we profiled developing Th2 cells using RNA-seq for gene expression, ATAC-seq (Assay for Transposase-Accessible Chromatin using sequencing) for chromatin accessibility and ChIP-seq of three key TFs: GATA3, IRF4 and BATF. We further acquired corresponding data from human donors to study the conservation of the regulatory pathway.

The regulatory function of all genes has been assessed by combining state of the art gene regulatory network analysis, comparison of Th2 *versus* Th0, early *versus* late, literature curation and genome-wide screen enrichment. Selected hits were validated in individual KO experiments. We characterize genes in terms of their impact on activation and differentiation, as well as on overall T helper cell identity. For ease of visualization, the integrated dataset is provided online at http://data.teichlab.org.

## Results and Discussion

### Genome-wide CRISPR/Cas9 screens recapitulate known genes and reveal novel genes in primary mouse T cell differentiation

Figure 1 depicts an overview of our experimental approach. First, a high complexity retroviral sgRNA library was generated (see online methods) (Figure 1b) and transduced into purified T cells from mouse spleens. We activated naive CD4+ T cells with anti-CD3 and anti-CD28 together with IL4 at day 0, using an optimized protocol to efficiently culture large numbers of primary cells. In essence, red blood cell lysis was used to quickly remove most cells, followed by magnetic bead selection. Culturing was then done on flat-bottom plates (see Methods for details). At day 1, T cells were transduced with the retroviral libraries and selected with puromycin from day 3. After dead cell removal, the screens were carried out on day 4.

**Figure 1:**
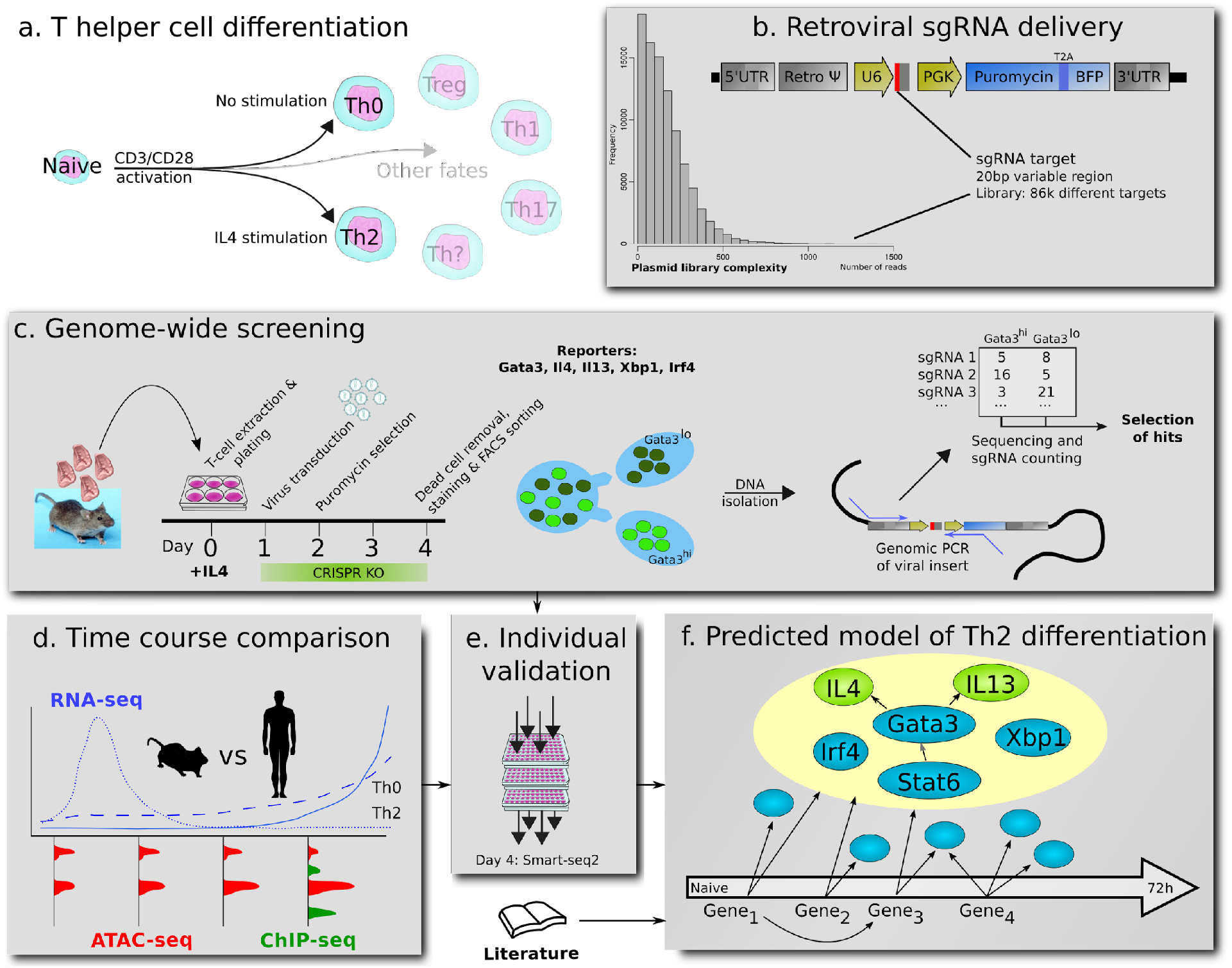
Overview of the experimental KO screening strategy. **(a)** In our culture system, naive, *ex vivo* T cells are differentiated into Th2 cells by IL4. Potential alternative T cell fates that may be open to genetically perturbed cells are indicated. *In vivo*, T cells develop into different subtypes dependent on stimuli. **(b)** The retrovirus is based on murine stem cell virus (MSCV), encoding one sgRNA per virus, and allows for BFP and puro selection. For the screening we have used a pool of plasmids, encoding over 86 000 sgRNAs, from all of which we produced viruses. **(c)** For genome-wide screens, we pool cells from up to 30 mice. After infection and puromycin selection, the cells are sorted based on fluorescence for the investigated gene. sgRNAs affecting gene expression are identified by genomic PCR. Differential sgRNA expression analysis then allows us to find genes affecting either viability (drop-out screen) or differentiation. **(d)** The top enriched and depleted genes (“hits”) were analyzed based on their dynamics measured by RNA-seq, ATAC-seq and ChIP-seq. **(e)** Particularly interesting genes were then further validated by individual KO and RNA-seq. **(f)** By using all this data and a curating the literature we provide a Th2 gene regulatory network.

Our screening strategy used two different approaches. For *Il4, Il13, Xbp1* and *Gata3*, we used T cells from transgenic mice carrying a fluorescent reporter driven by the promoter of the respective genes. In this protocol, cell populations with high or low fluorescence of the gene of interest were enriched with sgRNAs for upstream genes inhibiting or promoting Th2 cell differentiation, respectively. In addition, we carried out screens in which T cells were stained with antibodies for IRF4, XBP1 or GATA3. Most CRISPR screens to date are “drop-out” screens where the sgRNAs from an early time point are compared to those in the final surviving cell population. In contrast, here we identify differentiation-related genes by comparing the sgRNAs in the selected target-high *versus*-low fractions. We will refer to the most highly enriched or depleted genes (defined in more detail below) as “hits”.

In total, we carried out 11 genetic screens and analyzed them using the CRISPR screen hit calling software MAGeCK^28^. As an illustration of the results obtained, Figure 2a shows the hits in a screen using anti-Gata3 antibody staining (*i.e*. sgRNA for specific target genes), ranked by MAGeCK p-value, against the fold change (Th2, 0h *versus* 72h, described later) of those sgRNA targeted genes. Reassuringly, *Gata3* is recovered as a top hit in its own screen, as expected. Another top hit is a known signal transducer from the IL4-receptor to *Gata3*, the TF *Stat6*. Previous work has shown *Stat6* to be required for the majority of Th2 response genes in mouse and human^4,5^. This gives us confidence that relevant genes are recovered.

**Figure 2:**
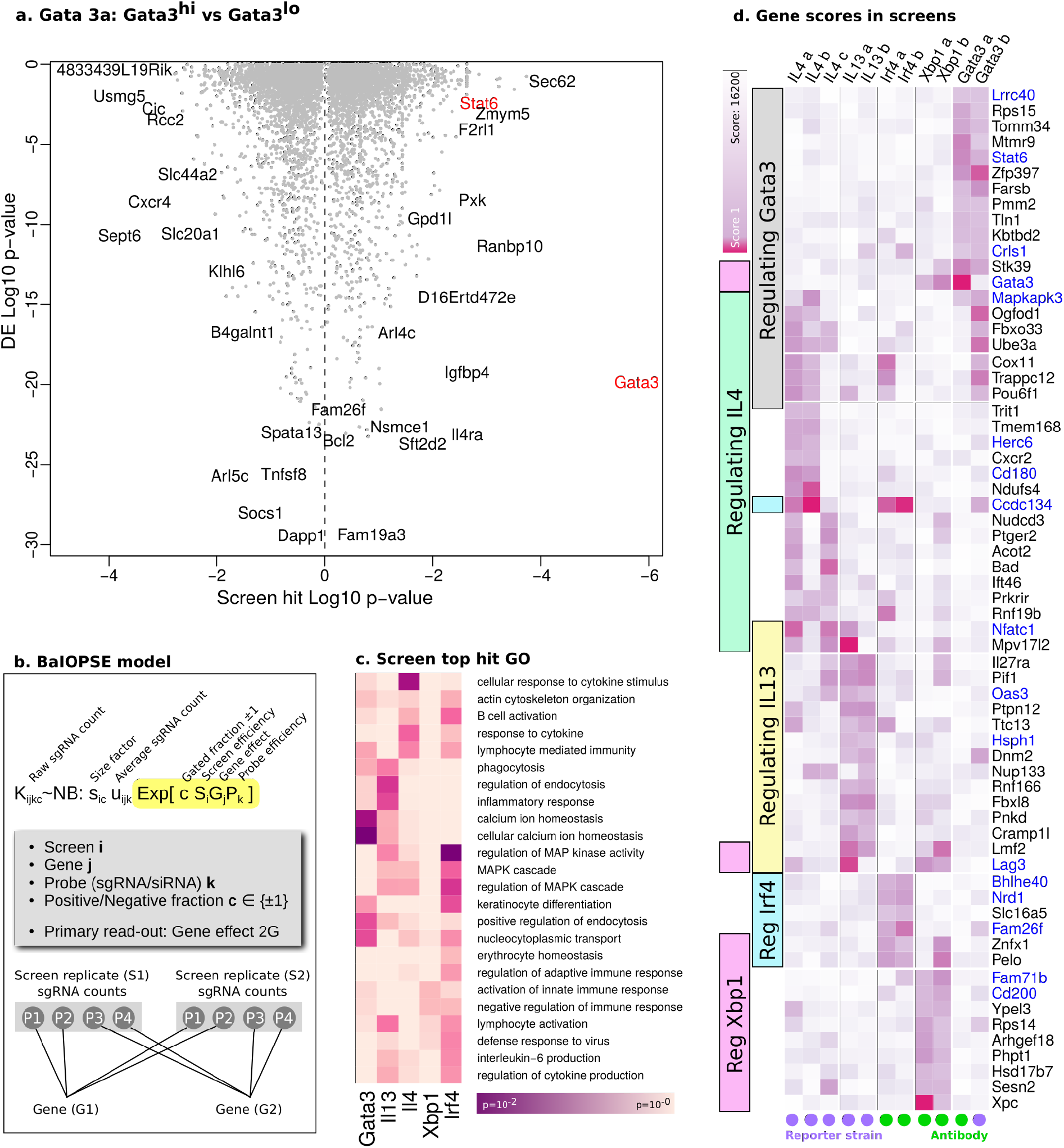
Results from genome-wide Th2 differentiation screen. **(a)** Hits from screen for *Gata3* expression measured by antibody staining. The x-axis denotes the p-value for differential expression obtained by MAGeCK (hits of high relevance toward both sides). The y-axis shows the p-value comparing Th2 and Th0 gene expression level (explained later). Highlighted in red are *Gata3* and *Stat6*, since these are known to control *Gata3* expression. **(b)** The alternative BaIOPSE (Bayesian Inference Of Pooled Screen Enrichment) hit calling model. This model is in essence an extended negative binomial differential expression model over sgRNA counts K. Each sgRNA has an efficiency term P, and each screen has an efficiency term S. The interesting read-out is the gene effect 2G. **(c)** GO annotation of top hits for each screen as defined by BaIOPSE. The color represents Log10 p-value. **(d)** Summary results of all 11 screens carried out. Genes that were consistent hits in multiple screens are shown (see methods for gene selection). The purple colour shows the Log10 combined MAGeCK rank (positive and negative enrichment combined). Screens that relied on antibody staining are marked by a green circle, and those based on fluorescent gene reporters are marked by a purple circle. Genes in blue have been KO:ed individually (see Figure 6).

As a further quality control we also compared the screens with an orthogonal hit calling model, BaIOPSE (Bayesian Inference Of Pooled Screen Enrichment, Figure 2b, further described in Supplementary methods). In short, size factors, screen efficiencies, probe efficiencies and gene KO effects are fitted simultaneously. Qualitatively, we find that there is reasonable overlap with MAGeCK^28^ and BaIOPSE (BaIOPSE scores in Supplementary File S13). In a GO analysis of top hits from all screens (Figure 2c), the categories for calcium and MAPK signaling have the lowest p-values. While BaIOPSE allows a more consistent integration of multiple screen replicates than MAGeCK, we use MAGeCK for the remainder of this paper because of its preexisting community acceptance and because BaIOPSE relies on informative priors.

In all subsequent descriptions of hits, we will refer to the expression of the targeted gene, rather than the level of sgRNA enrichment or depletion. We note that in Th2 cells (cultured 0h vs 72h) the fold-change in *Gata3* expression is relatively small, but still results in a strong phenotype. For the sake of brevity, in this paper we will use the nomenclature X^→y^, when gene X is in the top 5% of hits in the screen Y, either positively or negatively enriched. If gene X falls within the top 1% of ranked hits, we denote this as X^→y!^. A comprehensive list of all genes is included in Supplementary File S4 and results are summarised in Figure 2d.

We compared the hits with the immunological phenotypes of the International Mouse Phenotyping Consortium (IMPC)^29^. TFs with potential immune-related phenotypes, that were also hits in our screen, included the known T cell genes *Tcf7*^→Il4^ and *Xbp1*^→Il4^, but also several TFs not previously connected to T cells *e.g*. *Mbd1*^→Il13^, *Arid1b*^→Irf4^ and *Zfp408*^→Gata3^. *Mbd1*(−/−) mice develop autoimmunity, as *Mbd1* is important for *Aire* expression and early T cell development in the thymus^30^. The impact of *Mbd1* on *Il13* as shown here suggests an additional immunological role for this TF.

Next we identified hits that were consistent between screens (see Methods for details). Some genes appear to have a particularly strong impact on Th2 differentiation as they are hits in multiple screens. This includes the known genes *Il27ra*^→Il4,Il13!^ and *Lag3*^→Il4,Il13,Xbp1!^ but also genes not previously connected to T cells *e.g*. *Trappc12*^→Il4!,Ilf4!,Gata3!^, *Mpv17l2*^→Il4!,Il13!,Xbp1^ and the TF Pou6f1^→Il4!,Gata3^. The cytokine-like gene *Ccdc134*^→Il4!,Irf4!,Gata3^ is also a major hit. It has so far received little attention in the literature, but has been linked to arthritis^31^ and shown to promote CD8+ T cell effector functions^32^. In short, we have discovered many new genes with a broad effect on Th2 differentiation that deserve further investigation.

### Time-course analysis of gene expression and human-mouse comparison highlight metabolic genes

To place our hits into the context of Th2 development, we generated time-course data on mouse and human Th cells during both activation and differentiation (Figure 3a). Mouse and human primary Th cells were isolated from spleen and cord blood respectively and activated with anti-CD3 and CD28. The addition of IL4 to the medium resulted in the maturation into Th2 cells, while absence of IL4 resulted in activated “Th0” cells which proliferate but do not differentiate into a mature effector lineage (maturation here defined as the combination of activation and differentiation). We performed time-course bulk RNA-seq profiling on Th2 and Th0, and ATAC-seq at several time points during Th2 differentiation. The large number of data points allowed us to reconstruct the time-course trajectory of Th2 differentiation by principal component analysis (PCA), using RNA-seq data or ATAC-seq data alone (Figure 3b, c).

**Figure 3:**
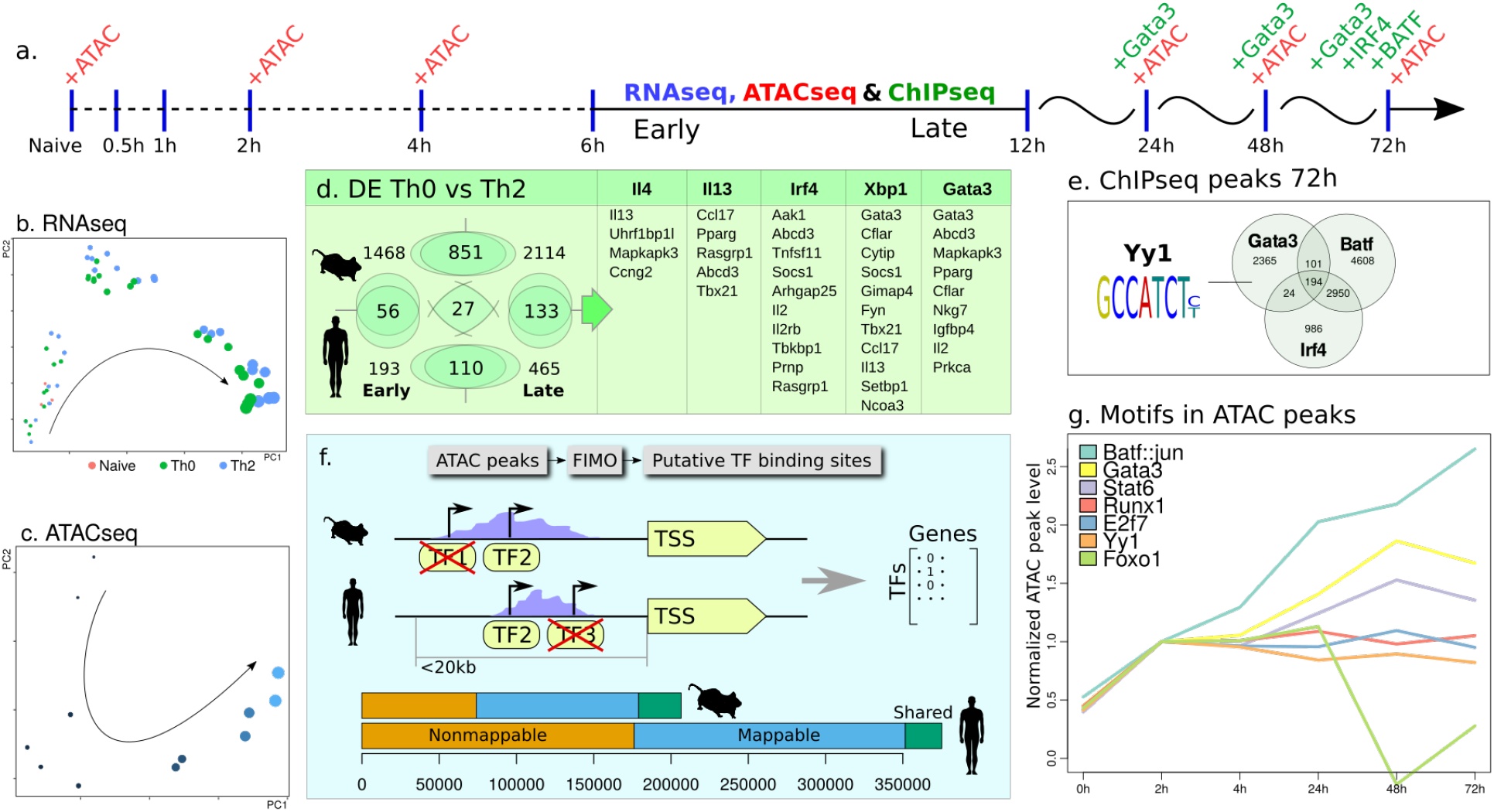
Molecular characterisation and assessment of hits over the time-course of Th2 differentiation. **(a)** The chosen time-points for RNA-seq, ATAC-seq and ChIP-seq. **(b)** PCA projection of bulk RNA-seq and **(c)** ATAC-seq samples. The size of the circle represents time. The naive samples separate in the third principal component not shown. **(d)** Number of differentially expressed genes in the early and late response, in human and mouse (p=10^−4^). DE genes in both human and mouse that are also hits in the genetic screens sorted by rank in their respective screen. **(e)** Workflow for finding conserved putative TF binding sites in human and mouse. The green region represents conserved (overlapping) peaks. The blue region represents peaks in regions with a corresponding sequence in the other species, but without peak conservation. The orange region depicts peaks lying in non-syntenic (unmappable) regions. **(f)** Examples of ATAC-seq peak dynamics associated with different TFs **(g)** ChIP-seq overlap and the YY1 motif found within the GATA3 peaks.

When carrying out differential gene expression (DE) analysis between the Th0 and Th2 populations, we split the time-course into an the early/fine-grained (0h-6h) period, and a late/coarse-grained period (0h + 6h-72h), as shown in Figure 3a. The number of DE genes is shown in Figure 3d, with the most DE genes also being hits in a screen highlighted. Importantly, a sizeable fraction of these (21%) were also identified in at least one of our genetic screens, providing orthogonal evidence for their importance (DE scores are in Supplementary File S14).

Evolutionary conservation supports functional relevance, so we carried out an equivalent RNA-seq analysis in cultured human primary T cells. Fewer DE genes were identified, possibly because genetic diversity between individuals may obscure some gene expression changes, but more than 1/5th of the human DE genes had direct orthologues in the mouse response. For the remainder of this paper we will refer to any gene being DE in either human or mouse, at any time, as simply DE.

A total of 216 genes were DE in both mouse and human, either early or late (*p*=10^−4^). DE genes that also are top hits in our CRISPR screens are shown in Figure 3d. We note the presence of the well-known cytokines *Ccl17*^→Il4,Il13,Xbp1^, *Il13*^→Il4,Xbp1^, *Il2*^→Irf4,Gata3^ and its receptor *Il2rb*^→Irf4^, and the TFs *Gata3*^→Xbp1!Gata3!^, *Tbx21*^→Il13,Xbp1^ and *Pparg*^→Il13,Gata3^. Several of these are canonical Th2 genes, but several novel hits were also found. Notably, several of these are related to metabolism, such as Pparg, which is thought to signal through mTOR and control fatty acid uptake^33^. Another metabolic gene, related to fatty acid transport^34^, with a strong phenotype in our screen, is *Abcd3*^→Il13,Ir4,Gata3!^ which has not yet been studied in T cells. The Th1-repressor *Mapkapk3*^→Il4,Gata3!^ is also a metabolic gene^35^.

Other hits have more diverse functions in T cell development. Hits include the known T cell regulator Stat-inhibitor *Socsl*^→Irf4,Xbp1^ (Suppressor of cytokine signaling 1). The *Il13* hit *Rasgrp1*^→Il13,Irf4^ (RAS guanyl releasing protein 1) is known to be involved in T cell maturation^36^ and links Guanyl to the RAS pathway. Interestingly another Guanylate protein, *Gbp4*^→Il13^, is also a novel Il13 hit (but with higher DE p-value). The novel *Il4* candidate regulator *Uhrflbp1*^→Il4^ has been connected to hypomyelination but could act through the chromatin regulator *Uhrf1* which is required for Treg maturation^37^.

In conclusion, a human-mouse comparison of DE genes highlights cytokines and TFs known to be important in Th2 differentiation, and suggests additional hits in our screens that are likely to be of functional importance, in particular novel genes that act as metabolic regulators (*e.g. Abcd3*).

### Analysis of chromatin dynamics reveals different TF binding patterns during maturation

To gain further insight into the regulation of gene expression we examined chromatin accessibility using ATAC-seq. The chromatin of naive T cells is condensed until activation. It has previously been shown that some TFs, for example *Stat5*, can only access the promoters of its target genes after T cell activation^43^. Th2 differentiation is classically thought to be driven by *Stat6* which in turn upregulates *Gata3*. We examined these dynamics over the time-course of the Th2 response.

The ATAC peaks were first called using MACS2^44^. Overall there is a massive gain of chromatin accessibility from 0 to 2h (Supplementary Figure 1 and Figure 3f). After this initial opening, the chromatin appears to recondense continuously and this process progresses past 72 hours, as indicated by the reduced total number of ATAC-seq peaks in each time point. We speculate that the regulatory network shifts from a general T cell network to subtype-specific network, and that cell identity becomes less plastic and less responsive to external perturbation over time.

We next compared TF binding predictions between human and mouse. Using FIMO^45^ we predicted TF binding sites within ATAC-seq peaks. To reduce the number of potential false positive peaks we concentrated on ATAC peaks that are conserved between mouse and human by calculating the percentage of overlapping peaks between species (10–15%) (Figure 3e) and used these conserved binding sites for the rest of the analysis.

For different TFs, we examined how ATAC peaks, in which the relevant TF motif is found, are changing over time (Figure 3g). Peaks containing the previously mentioned YY1 motif are stable or decrease slightly. This finding, together with the strong phenotype in our screen, reiterates Yy1 as a key supporter, but not driver, of Th2 differentiation. BATF::JUN is one of the (composite) motifs associated with the largest increase in relative peak size. This is consistent with the suggestion that BATF can act as a pioneer factor to open chromatin11. Of the TFs identified in the screen, *Jun*^→Il13^, *Fos*^→Irf4,Xbp1^, *Fosl2*^→Gata3!^ and *Gata3*^→Xbp1,Gata3^ are all associated with increasing peak height. Since Jun/Fos and Fosl2 all recognize the same AP-1 motif, the exact TF composition at these peaks is likely to depend on their expression level. Notably, *Fosl2* expression is highest at 1–2h in Th0/2 with largely unchanging levels in Th1/2/17/Treg^46^. Overexpression of *Fosl2* has been shown to block IL17A production in Th17 by competing for AP1-sites^11^, but overall *Fosl2* expression is low in lymphoid cells^42^. *Fos*, *Jun and Jund* are also transiently expressed during the first 6 hours. *Jund* is the only classical AP-1 factor that has increased expression over time albeit slowly. As most AP-1 factors are expressed at low levels, we speculate that *Batf*, whose expression increases continuously, is the primary driver behind these peaks.

At the other extreme, some TF motifs are overrepresented in peaks that decrease over time, such as *Hoxd9*^→Il4^, *Atf3*^→Gata3^, *Atf4*^→Il4!^, *Foxj2*^→Gata3^, *Dmbx1*^→Irf4^, *Foxa2*^→Il4!^, *Foxo3*^→Il4^ and *Foxc2*^→Il13!^. Several of these TFs also have low or decreasing expression levels. We have previously shown that *Atf3*^→Gata3^ positively regulates *Ifng*^47^ and promotes Th1 differentiation in humans. *Atf4*^→Il4!^ has been shown to be important for Th1 function as stress regulator^48^ but the impact on *Il4* extends this claim to Th2. *Foxo1*^→Il13!Xbp1!^ is a highly expressed TF but peaks containing this motif are also decaying. *Foxo1* has recently been shown to inhibit H3K27me3 deposition at pro-memory T-cell genes^49^. *Foxj2* has similar behaviour to *Foxo1* but has not been studied in T cells. However, a link has been made between *Stat6, Foxj2* and cholesterol in lung cancer cell lines^50^. Because of the importance of *Stat6* in Th2 such a connection would also be interesting in T cells. Inferred STAT6 binding sites were also compared with previous mouse and human data^4,5^, and we found that the vast majority of the previous target genes are also DE in our time course analysis (Supplementary File S10, S11). A list of all TFs and the average height of peaks containing their cognate motif is provided in Supplementary File S7.

To further characterise the dynamics of the Th2 response, we generated ChIP-seq data at several time points (Figure 3a) for the known Th2 TF GATA3, as well as for the TFs BATF and IRF4 which we found to be involved in increasing ATAC peaks. We created a mouse strain with a 3xFLAG-mCherry GATA3 construct (T2A fusion) for this purpose (see Methods for details). The ChIP-seq peaks for *Batf* and *Irf4* have a large overlap as previously reported^11^ (Figure 3e) (Jaccard index=0.35). However, we saw no significant overlap of these two factors with GATA3 (Jaccard index=0.028 and 0.032), suggesting that any collaboration between GATA3 and BATF/IRF4 is not due to direct co-localization on the chromatin. MEME^38^ was applied to the sequences in the GATA3 peaks to find other potential binding partners, and we found enrichment of YY1^→Il4,Il13,Xbp1,Gata3^ (*p*=2.5*10^−58^) binding motifs. This is consistent with previous reports that *Yy1* overexpression is sufficient to drive Th2 cytokine expression^39^.

Focusing on GATA3 with its 10,203 peaks, a GO-term analysis of its nearby genes yielded “natural killer cell activation” (*p*=6*10^−3^), but included few other immune-related terms. This is likely to be due to the fact that *Gata3* has distinct roles in other cell types^40,41^ (a survey has shown that its expression is highest in breast cancer cell lines^42^). However, since we performed time-course ChIP-seq we were able to selectively investigate peaks based on their dynamics. Genes near peaks decreasing over time were not linked to any particular immune-related GO-terms, but a GO term enrichment for genes near increasing peaks revealed “defense to bacterium”/”viral life cycle” (*p*=5*10^−3^) as the top term, and included other terms such as “myeloid leukocyte activation” (*p*=2*10^−2^). From this data we speculate that using time-course ChIP-seq we can split the GATA3 targets into those where GATA3 has a passive role (e.g. priming the chromatin for other TFs), and targets where GATA3 is driving the changes. A ranking of peaks and nearby genes, as well as GO terms, are provided in Supplementary File S9.

To conclude, the early change in ATAC-seq peak size over time reflects a rapid increase in accessibility for all TFs. In addition to the global change we find pioneer factors driving the opening of chromatin and we find TFs whose binding is reduced with time. Amongst our screen hits we find genes in both categories. These are all likely to be functionally important as either activators or repressors for the specific T helper type.

### Correlation analysis find critical paths in the regulatory network

So far, hits have been considered one at a time. The generation of transcriptional networks potentially allows us to integrate the effect of multiple regulators. A regulatory network was created using ARACNe-AP^51^ on the mouse time-course RNA-seq data (full network in Supplementary File S12). In essence, edges were created between genes with high mutual information (correlation in gene expression datasets). In this network the screen hits cluster together, in particular, those for *Il4* (p=10^−2^ for combined rank<300), but also those for *Gata3, Il13* and *Irf4*. The upstream genes of *Xbp1*^→Il4^ have a similar, but weaker trend. Hits from one screen also cluster together with hits from other screens, suggesting functional overlap.

The full network is too complex to be visualized. However individual regulons with a high number of hits are displayed in Figure 4a, filtered by ATAC conserved binding sites. This filtering also imposes a directionality on the edges. As an example we depict the *Nfkb2* regulon which encompasses hits for the *Il13, Il4, Gata3, Xbp1* and *Irf4* screens. For each gene enriched in a particular screen, a binomial test was performed to see if the connected genes are also enriched in the screen. For the ATAC-filtered time-course network of DE mouse genes, *Ube2m*^→Il4!Gata3!^ has 4 out of 16 genes affecting *Il4* (p=5*10^−3^). Only *Ybx1*^→Il4^ (*p*=1.1*10^−2^) has a similar level of *Il4* enrichment. *Irf8*^→Xbp1^ is the regulon most enriched for hits from the *Xbp1* screen (p=10^−3^), followed by *Hkl*^→Il4,Xbp1^ (p=2*10^−3^) and *Sqstml*^→Xbp1!^ (p=10^−2^). The regulon most enriched for *Irf4-hits* is the Arginyl aminopeptidase, *Rnpep*^→Il4,Irf4,Gata3^ (*p*=4*10^−2^). *Rnpep* is regulated by a number of genes that are hits from different screens and *Rnpep* itself is a regulator for *Il4, Irf4* and *Gata3*, suggesting a commonly involved pathway. This gene has similarities to other aminopeptidases, and drugs inhibiting these have been shown to modulate the immune response^52^. *Rnpep* thus represents an alternative and potent entry point for these drugs.

**Figure 4:**
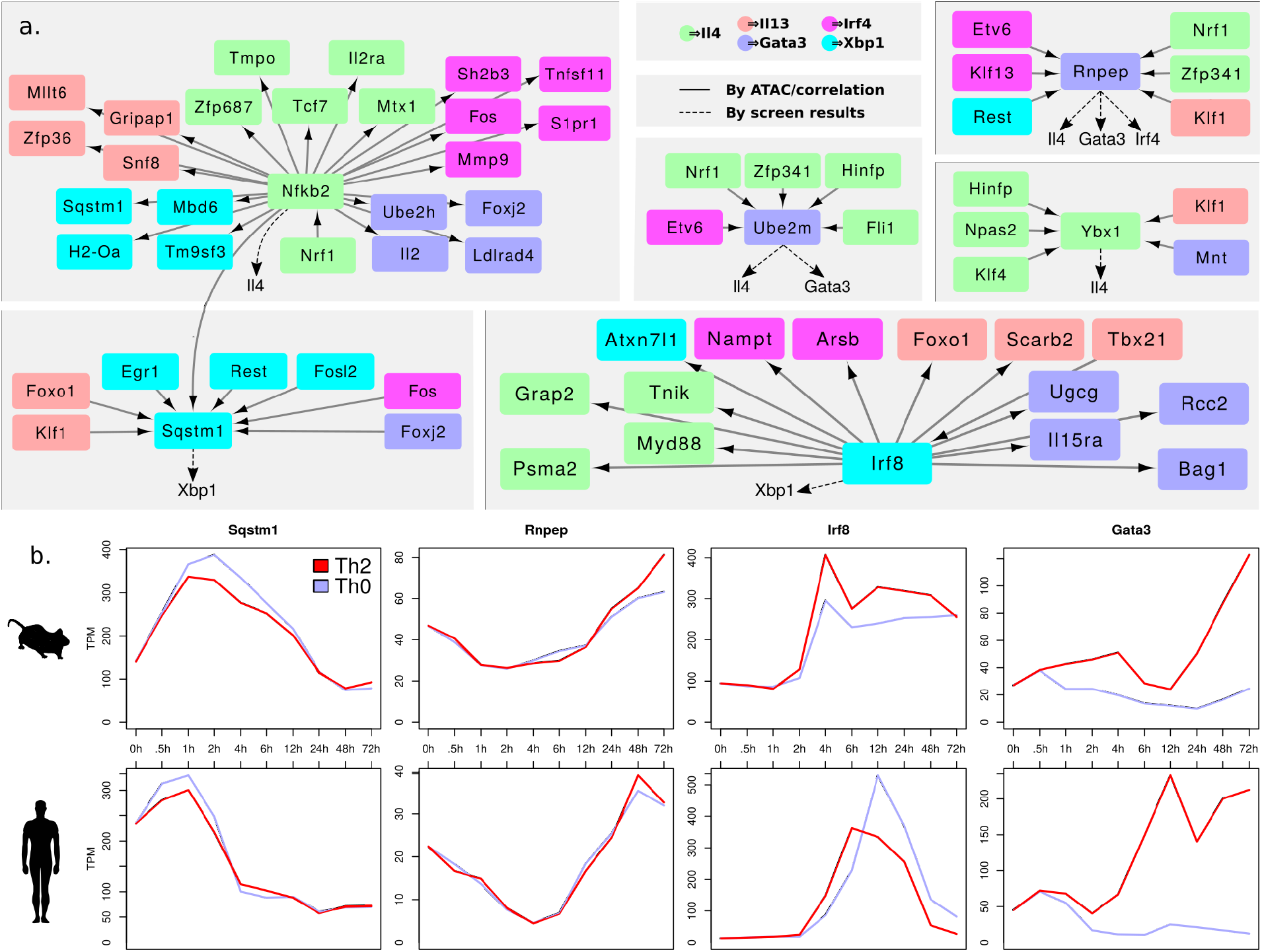
Correlation analysis finds critical paths in the regulatory network. **(a)** Examples of regulons from the ARACNe network of time-course RNA-seq with high enrichment in hits. Hits from different genetic screens are shown in colour as indicated. Dotted arrows highlight how some genes, e.g. *Rnpep*, may act in commonly involved pathways **(b)** Gene expression over time for some of the genes highlighted by ARACNe. *Gata3* is included as a reference point.

The levels of the above mentioned genes were also compared between different Th types^46^. *Ube2m* was specific to iTreg, *Sqstm1* to Treg/Th17, and *Irf8* and *Tbx21* had variable expression.

*Rnpep* was slightly elevated in Th2 vs Th0 but only weakly DE. The examples we have highlighted were not DE between Th2 and Th0 (Figure 4b). This highlights the orthogonality of the ARACNe-based network approach and our ability to extract hits which are neither DE nor TFs, but nevertheless play an important regulatory role in Th2 differentiation.

### Motif activity analysis quantifies transcription factors controlling activation *versus* differentiation

The ARACNe approach is based on similarities in gene expression between a given TF and its potential targets. However, TF function is frequently determined post-transcriptionally, for example, STAT6^→Gata3^ activity is regulated by phosphorylation, and TBX21^Xbp1,Il13^ in part operates by sequestering and displacing GATA3^→Gata3,Xbp1^ once expressed^10^ (100-fold up-regulated in Th1^46^). In contrast, the ISMARA^53^ algorithm builds a network by linking TFs to potential target genes based on the presence of the relevant motif in an ATAC-seq peak within the vicinity of the transcription start site (TSS) of that target gene. Edges are then curated by searching for co-regulation amongst predicted targets of a given TF (see methods). Interestingly we find that there is very high correlation in the predicted network when comparing results obtained with sites predicted from ATAC-seq peaks to results obtained with measured TF binding (ChIP-seq data) (Figure 5b), suggesting that the algorithm performs well on ATAC-seq input data, allowing us to analyse all TFs, not just those with ChIP-seq data.

**Figure 5:**
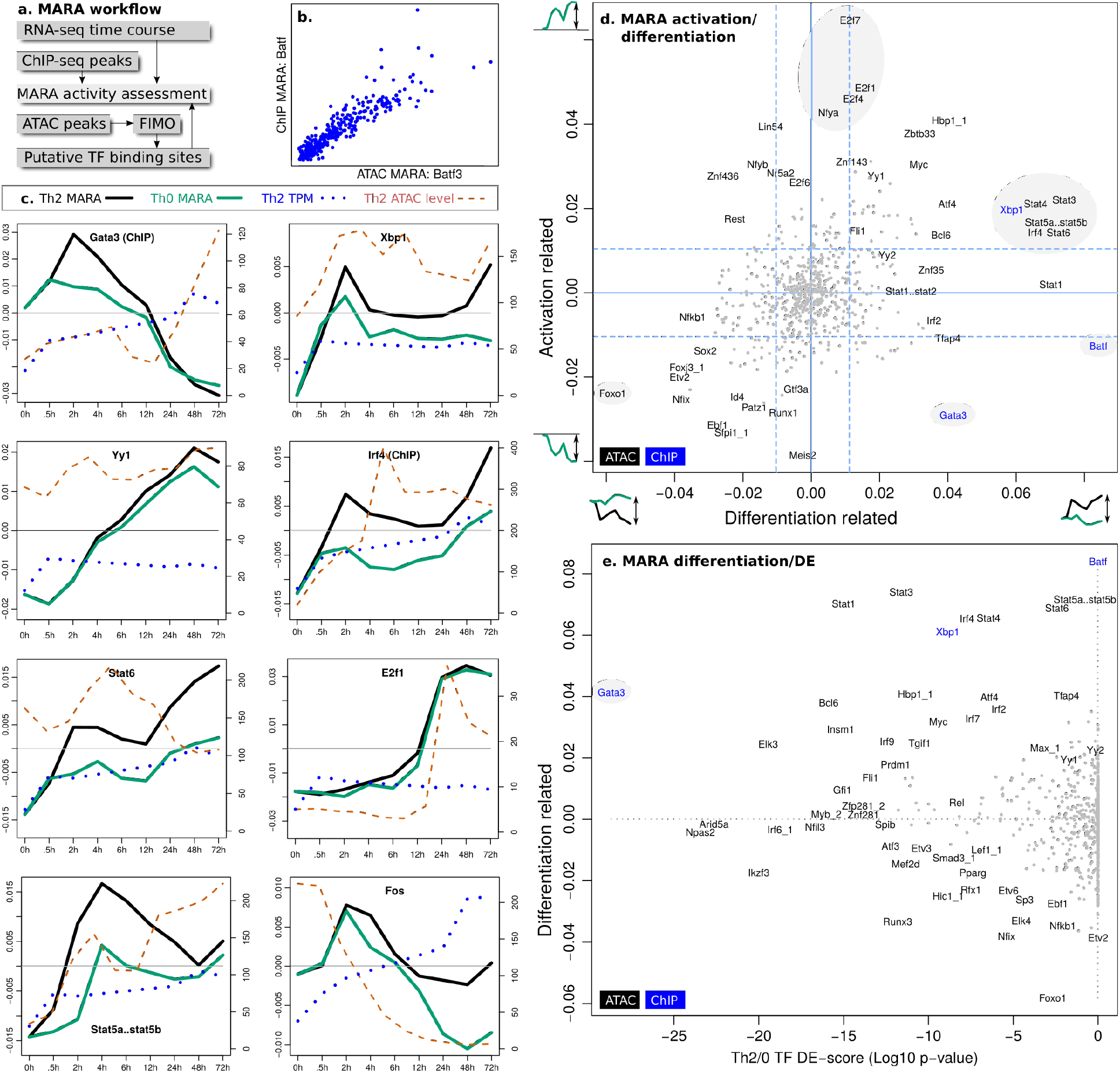
Analysis of transcription factor activity using “Motif Activity Response Analysis” (MARA) **(a)** Workflow for combining putative binding sites with time-course RNA-seq. **(b)** Comparison of BATF activity predictions for individual genes, by ATAC-seq predicted binding sites and ChIP-seq peaks. **(c)** Dynamics of selected TFs, comparing their expression level, activity in Th2 (black line) and Th0 (green line) and chromatin accessibility. **(d)** MARA activation vs differentiation scores (as defined in text) of all TFs. **(e)** Comparison of differentiation score and DE *p*-value Th2 vs Th0.

To obtain an overview of the role of all the TFs we have categorized factors according to their activity over time within the Th2 differentiation pathway, and whether their activity differs when comparing Th2 to Th0 cells. In other words, two distinct comparisons are made: Firstly t=0h *versus* t=72h within the Th0 compartment, which we term “activation”, and secondly Th0 *versus* Th2 cells at t=72h, which we term “differentiation”. Figure 5c illustrates this analysis by showing MARA activity scores independently calculated for Th2 and Th0 cells for a number of selected TFs. An example of a TF strongly associated with differentiation (i.e. large difference between black and green lines) is *Fos*^→Irf4,Xbp1^, while an activation phenotype, reflected in a large difference between t=0h and t=72h, is observed for *E2f7*^→Il13^. The majority of TFs display a behaviour reflecting both activation and differentiation (Figure 5d). The calculated activity score was also related to expression highlighting that for many TFs activity does not closely correlate with expression (Figure 5c, e). The Th2-defining TF *Gata3*^→Gata3!,Xbp1!^ shows transient activity but its expression increases with time. *Gata3* is one of the strongest mediators of both activation and differentiation, although its differentiation activity appears to be exerted early. *Stat6*^→Gata3!^ is also thought to act early in differentiation, after its activation via the Il4 receptor *Il4ra*^→Gata3^. We previously showed that during Th2 differentiation, signals from IL4R are predominantly transduced through STAT6^5^. Consistent with those findings, our data suggests that *Stat6* activity continues to increase throughout differentiation. Interestingly all the STAT proteins map closely together in Figure 5d, affecting primarily differentiation but also activation, possibly all contributing to different extents depending on their expression, phosphorylation status and interactions with other proteins and regulatory elements. *Irf4* is also in this cluster (Figure 5e). *Foxo1*^→Il13,Xbp1^ and *Xbp1*^→Il4^ are also strongly connected to activation and differentiation, but with *Foxo1* in the opposite direction. Previous work suggested that the primary role of *Batf* is to open the chromatin together with *Irf4*^11^, and this is consistent with our analysis in Figure 3g. Here it is one of the strongest differentiators, suggesting that chromatin opening is restricted to sites required for differentiation. The roles of other genes are more diffuse. TFs that were identified as hits include *Atf4*^→Il4!,48^ and Yy2^→Gata3^, *Id4*^→Il13(II4),Xbp1^, *Ebf1*^→Irf4^, *Foxp2*^→Gata3^, *Yy1*^→Il4,Il13,Xbp1,Gata3^ and *Fli1*^→Il4!^ affecting both activation and differentiation, but with weaker effects. The identification of a cluster of E2F-proteins as strongly and purely activation-related is consistent with their role in cell cycle control.

The MARA approach allowed us to extract canonical Th2 TFs, such as *Stat6, Gata3* and *Batf*, and in addition highlighted newly identified TF-hits (*E2f7, Foxo1*) likely to be relevant for Th2 activation and differentiation. Since MARA is not directly dependent TF-target gene co-variation, the output is complementary to the previous DE and ARACNe approaches.

### Validation of hits by individual CRISPR KO assess gene influence on activation vs differentiation

Next we used the results described so far, related this to the existing literature, and chose a panel of 45 genes (40 by screen scores), which were then validated by individual CRISPR KO. Several of the chosen genes have been studied before though not specifically in T cells. Our selection of interesting genes for further characterization is by no means comprehensive, and additional genes can be found by browsing our online resource.

For each KO, cells were grown under Th2 differentiation conditions and RNAseq carried out on day 4. For each gene a DE list of KO vs non-targeting control was derived and compared to the activation and differentiation axes (Figure 6a). We defined the activation axis as the DE genes from Th0(0h) vs Th0(72h) and the differentiation axis as the DE genes from Th0(72h) vs Th2(72h) (Figure 3a,b). It should be noted that some genes might not be consistently higher or lower in Th2 vs Th0 cells. There are thus many other possible definitions of the differentiation axis. To find if a KO aligns with one of these axes, the DE genes of the KO are compared to the DE genes of the axis (see Supplementary methods). Figure 6a shows that all genes tested map away from the neutral centre of the plot (shaded in grey), indicating that the hits contributed to either differentiation or activation, but mostly both, thereby validating the relevance of these genes in Th2 maturation. In this analysis, *Il4* however shows little effect but we believe this is due to IL4 being supplemented in the media. Consistent with the MARA analysis, *Stat6* is primarily differentiation related. By basing this analysis just on expression, GATA3 now appears to be primarily activation dependent. However, for TFs the MARA analysis accounts for activity and therefore likely to be more accurate if the two analyses diverge. The majority of KO genes affect both differentiation and activation. Examples of genes which have not been studied extensively before in T cells are *Pgk1*^→Il4^, *Lrrc40*^→Gata3^, *Slc25a3*^→Irf4^ and *Ccdc134*^→Il4!,Irf4!^.

**Figure 6:**
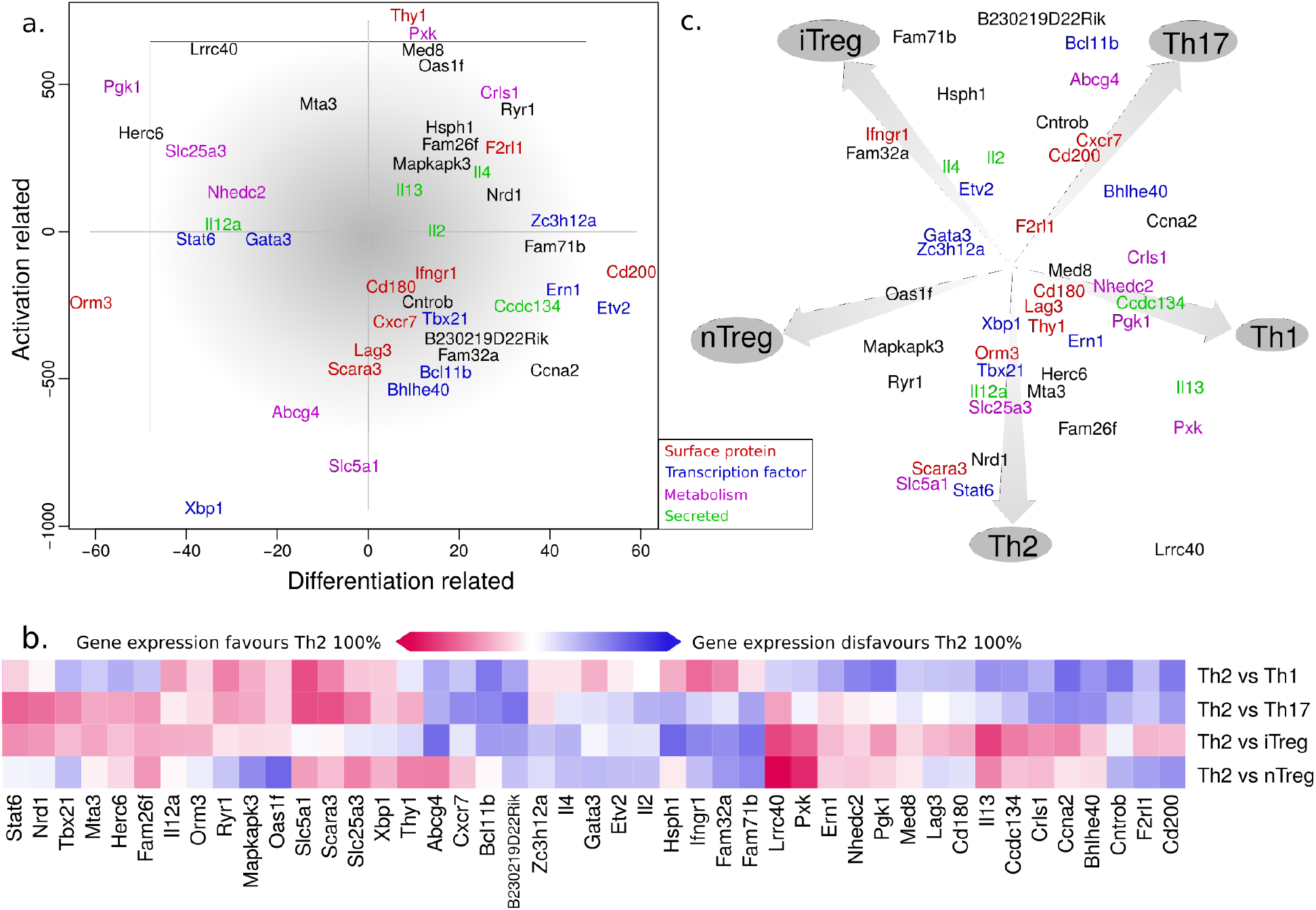
KO effect on T helper cell identity. **(a)** The effect on differentiation and activation of gene KOs. DE genes between KO and WT are identified and compared to DE genes lists defined for activation Th0 (t=0) vs (t=72h) and Th2 vs Th0 (t=72h) (further defined in text). Genes/KOs toward the middle of the plot have the least effect. **(b)** Gene influence on Th identity. The KO DE gene lists are compared with fate DE list, both quantitatively (% overlap, shown by colour intensity) and qualitatively (same or opposite direction, depicted in blue and red). **(c)** A dimensionality reduced summary of 6b. Note that a central position does not reflect a Th0 phenotype since all comparisons are relative to the IL4 induced Th2 state. It reflects either very small changes from Th2, or opposing effects in different T fates that can cancel each other out in this graphic representation.

### Individual CRISPR KOs reveal genes affecting multiple Th programs

The activity/differentiation analysis only considers Th0 vs Th2. However extensive previous work indicates that different Th fates are interrelated and gene KOs may push cells towards different fates. To assess this, we first developed a DE list of KO *versus* WT genes. Next, using previously published data46, we compared day 5 gene expression signature of Th2 to that of other culture conditions (Th1, Th17, iTreg and nTreg), generating a second DE list. The overlap of the two lists is assessed (Figure 6b, Supplementary Methods) by amount (%, shown by colour intensity) and direction (same or opposite, depicted in blue and red).

A complex pattern emerges that further emphasises the complexity and plasticity within the T helper cell compartment. A simplified overview using a dimensionality reduction is also provided (Figure 6c). In this visualisation, all data points start in the middle and are then moved towards or from different fates by vectors representing the similarity between the KO DE list and the fate DE list, as indicated by colours in Figure 6b. By design, KOs having a small or ambiguous effect are placed in the middle. In contrast, high *Stat6* expression pushes cells towards a Th2 phenotype in three of the four different comparisons, with the effect that Stat6 maps very closely to the Th2 fate in Figure 6c. We note that a central position does not reflect a Th0 phenotype since all comparisons are relative to the IL4 induced Th2 state. It reflects either very small changes from Th2, or opposing effects in different T fates that can cancel each other out in this graphic representation.

Our analysis illustrates that the different T cell fates are connected, but that their relationships are complex. Some genes can favour both Th17 and Th1 (*e.g. Bcl11b*^→Il13!^), or Th17 and iTreg (e.g *Hsph1*^→Il13!^), or nTreg and iTreg (e.g *Fam32a*^→Gata3^) or lastly, nTreg and Th1 (*e.g. Cd200*^→Xbp1^) (Figure 6b). Inspecting individual genes, *Stat6*^→Gata3^ is shown to promote Th2, and *IL13*^→Il4,Xbp1^ is identified as pro-Th1. *Il13* is known to be dispensable for Th2 and can, in fact, be produced by Th1 and Th17^54^. The dimensionality reduction pinpoints additional strong pro-Th2 gene hits: *Nrd1*/*Nrdc*^→Irf4^, which has recently been shown to regulate inflammation in the murine stomach by controlling *Tnfa* shedding^55^, and *Slc5a1*^→Ifr4^ (glucose/galactose transport). Further Th1/2-regulators might include *Slc25a3*^→Ifr4^ (transporter of phosphate into mitochondria), *Herc6*^→Il4^ (E3 ligase for ISG15, important for antiviral activity^56^), the elusive^57^ *Thy1, Pgk1*^→Il4^ (glycolysis-related enzyme) and the novel *Lrrc40*^→Gata3^. Possibly the most interesting gene is the secreted *Ccdc134*^→Il4!,Irf4!,Gata3^ mentioned in a previous section. Here we show that this potential cytokine favours a Th1/2/17 fate over the Treg state.

In summary, we have carried out a gene-by-gene validation for a number of selected Th2 regulators. RNA-seq patterns for KOs of previously known regulators, such as *Gata3* and *Stat6*, are consistent with their ascribed role. However, in addition to this, we find that some of the newly identified hits demonstrate extremely interesting phenotypes with respect to T helper differentiation. These are exemplified by *Abcg4*^→Irf4^, *Fam32a*^→Gata3^, *Slc5a1*^→Irf4^, *Herc6*^→Il4^, *Ccdc134*^→Il4!,Ilf4!,Gata3^, *Lrrc4CT*^Gata3^ and *Nrd1*^Ifr4^, all of which deserve further study.

## Conclusions

In this study, we demonstrated, for the first time, the applicability of CRISPR to primary murine T cells. By carrying out *in vitro* genome-wide screens we have created a resource of genes important for T helper cell differentiation. We provide optimized protocols for performing additional screens as well as individual KOs. In our hands, these methods have not only worked better than RNA interference, but CRISPR also has advantages in terms of improved targeting and gene disruption instead of down-regulation. In our analysis, we have chosen 5 different read-outs *(Gata3, Il4, Il13, Xbp1* and *Irf4*), each known to be associated with Th2 differentiation. The fact that the different screens we have carried out generated many overlapping hits increases our confidence that relevant phenotypes were chosen.

In addition, we generated RNA-seq data for both mouse and human T cells following a time-course of Th2 cell differentiation. We provide a website (http://data.teichlab.org) that allows one to query individual genes with respect to Th2 expression dynamics and chromatin accessibility. This also includes the corresponding human time-course and plasticity analysis. By combining our CRISPR KO screen with time-course data, we have been able to provide a comprehensive map of the most important genes for Th2 polarization.

Key to our study is a systematic and unbiased approach to interrogate genes that contribute to Th2 differentiation. The hits we identify belong to many different classes of proteins, including cytokines, TFs, proteins involved in calcium signalling and metabolic genes. The precise mechanisms by which they impinge on the regulatory network is still unclear in many cases, but an exception are transcription factors. We know these regulate large numbers of genes by virtue of binding to their promoters as well as enhancers, and protein-DNA interaction can be profiled genome-wide through the time-course of differentiation. We have taken advantage of this in our study and have employed RNA-seq, ATAC-seq and ChIP-seq to dissect the gene regulatory network. This has allowed to us to identify TFs that play an important role in the development and have also been highlighted by our genetic screens.

We interpret our findings in the context of “Waddington’s landscape” (Figure 7a) which can be applied to any gene. Within the developmental landscape of Th2 differentiation, we distinguish between two modes of action: activation and differentiation. Genes related to activation either drive cells towards or inhibit cells from entering the cell cycle and the Th0 state, which we quantify as the difference between early and late gene expression. Genes related to differentiation are selected based on the difference between Th2 and Th0. Genes are sorted according to these two axes in Figure 7b. Notably, only a few factors are purely associated with either activation or differentiation, and the vast majority are involved in both processes (Figure 6a). This suggests pervasive intertwining of the regulation of activation and cell cycle entry on the one hand, and differentiation on the other hand.

**Figure 7:**
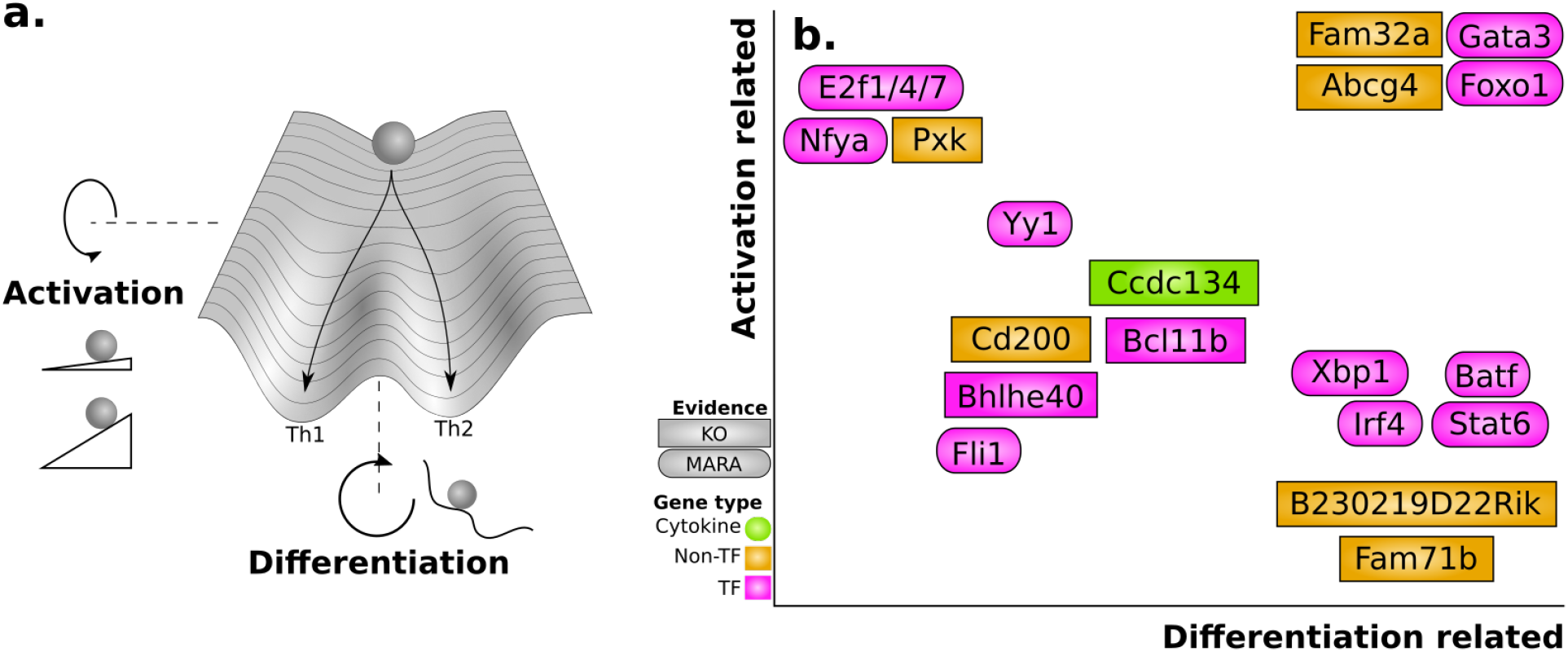
A conceptual view of Th2 differentiation. **(a)** While the genes controlling Th2 fate are from a wide range of programs, their behaviour can be categorized into the modes of activation and differentiation. Rotation of the landscape around the back to front or left to right axis will enhance differentiation and activation respectively. **(b)** Some of the genes according to these categories, including several genes identified in our screens that were not previously characterised in a T cell context.

Our analysis of Th2 differentiation has allowed us to re-discover known regulators, such as *Gata3, Stat6* and many others. In addition, we have highlighted a number of novel or only poorly studied genes that impact Th2 cell formation. Amongst TFs, examples include *Foxo1*^→Il13,Xbp1^, *Bcl11b*^→Il13!^, and *Bhlhe40*^→Ifr4^ (Figure 6b). Non-TFs have also been highlighted, including the cytokine *Ccdc134*^→Il4!,Irf4!,Gata3^.

For 45 genes, we also generated specific knock-outs, phenotyped the resulting cells and compared their gene expression patterns to that of different T helper subsets. We observe highly variable patterns for different gene knock-outs. We believe that the complexity of these patterns reflects the interrelatedness of different T cell subsets. In other words, we find that a gene KO can change the phenotype of a cell towards gene expression patterns reminiscent of more than one subtype, for example, both Th1 and Th17 or both Tregs and Th17 and so on. This is consistent with the idea that during the early stages of differentiation (0–72 h) the barrier between different fates is low and can be shifted by a change in expression of a single factor.

Our results yield a list of genes involved in T helper cell differentiation that deserve further analysis, and an efficient protocol for CRISPR-mediated KO. Both of these are key tools that will enable a more complete understanding of T helper cell biology.

## Methods

See on-line methods for further information

## Author contributions

R.M. helped with cell culturing and early trials of virus production and infections. X.C. and U.U. designed and performed the time-course RNA-seq, ATAC-seq and ChIP-seq. X.C. further helped with western blots, antibody staining and DNA isolation from fixed cells. T.G. performed the time-course data spline DE and PCA. K.M. advised on the ARACNe analysis and helped to write the manuscript. G.D. derived the ES cells from the GATA3-FLAG mouse. K.Y. provided the *Cas9* mice, the sequencing protocol and advised on cloning. R.L. and S.T. developed the experimental design for the RNA-seq, ChIP-seq and ATAC-seq time course for human and mouse cells. R.L. and S.T. initiated and supervised the study. S.T. contributed to the experimental design, data interpretation and writing of the manuscript. J.P. helping with T cell culturing, antibody staining and mouse maintenance. J.H. did the remaining work, in particular, the cloning, the CRISPR screening, the individual CRISPR KO, analysis and wrote most of the paper.

## Acknowledgements

We would like to thank Natalia Kunowska, Xin Xie, Andrew Knights and Sebastian Łukasiak for discussions about 293T culturing, CRISPR and virus production. Bee Ling Ng, Chris Hall and Jennie Graham helped with cell sorting. We thank all voluntary blood donors and personnel of Turku University Hospital, Department of Obstetrics and Gynaecology, Maternity Ward (Hospital district of Southwest Finland) for the cord blood collection. We thank Marjo Hakkarainen and Sarita Heinonen for their technical assistance.

## Data and materials availability

All the plasmids including the plasmid library are available from Addgene (see Online Methods for accession numbers). Selected parts of the data are also available for online visualization at http://data.teichlab.org. The sequencing data has been deposited at ArrayExpress (E-MTAB-6276, E-MTAB-6285, E-MTAB-6292 and E-MTAB-6300) and the R code used for the analysis is available on Github (https://github.com/mahogny/th2crispr).

## Supplementary files

S1. Supplementary figures

S2. Oligos for creating targeted sgRNA plasmids

S3. Primers for qPCR and cloning

S4. Screen sgRNA counts and MAGeCK aggregate scores

S5. Raw count and condition tables for targeted KO RNA-seq

S6. DESeq2 output for follow-up KO and RNA-seq

S7. TF-chromatin associations over time

S8. MARA input and output files

S9. Time-course behaviour and GO-analysis of GATA3-associated peaks

S10. Overlap with previous human study of Il4/Stat6-targets

S11. Overlap with previous mouse study of *Il4*/*Stat6*-targets

S12. ARACne network

S13. BaIOPSE screen scores

S14. RNA-seq time course TPM and DE scores

## Online methods

### Ethics statement

The mice were maintained under specific pathogen-free conditions at the Wellcome Trust Genome Campus Research Support Facility (Cambridge, UK). These animal facilities are approved by and registered with the UK Home Office. All procedures were in accordance with the Animals (Scientific Procedures) Act 1986. The protocols were approved by the Animal Welfare and Ethical Review Body of the Wellcome Trust Genome Campus. The usage of the cord blood of unknown donors was approved by the Ethics Committee of Hospital District of Southwest Finland.

### Cloning

The software Collagene (http://www.collagene.org/) was used to design and support the cloning. Phusion polymerase (NEB #M0531L) was used for all cloning PCR reactions.

The entire BFP/puromycin and sgRNA system was PCR-amplified from pKLV-U6sgRNA(BbsI)-PGKpuro2ABFP (primers: kosuke_mfei_fwd/kosuke_clai_rev). The plasmid pMSCV-IRES-mCherry FP (Addgene #52114) grown in dam-/dcm-competent *E. coli* (NEB #C2925I), was digested with NEB ClaI/MfeI and the backbone was gel purified using the QIAquick Gel Extraction Kit (Qiagen #28704). Ligation was done with T4 ligase (NEB #M0202T). The resulting plasmid that can be used to target individual genes was named pMSCV-U6sgRNA(BbsI)-PGKpuro2ABFP (Addgene #102796).

To produce the pooled library pMSCV-U6gRNA(lib)-PGKpuroT2ABFP (Addgene: #104861) the sgRNA part of a previous mouse KO sgRNA pooled library22 (Addgene: #67988) was PCR-amplified using the primers gib_sgRNAlib_fwd/rev. Up to 1ug was loaded in a reaction and run for 10 cycles. The insert was gel purified, and then repurified/concentrated using the MinElute PCR Purification Kit (Qiagen #28006). The backbone from pMSCV-U6sgRNA(BbsI)-PGKpuro2ABFP was obtained by BamHI-HF (NEB) digestion. The final product was produced by Gibson assembly (NEB #E2611S) and combining the output of 10 reactions. 6 tubes of 5-alpha Electrocompetent *E. coli* (NEB #C2989K) were transformed using electroporation and the final library obtained by combining 4 maxipreps. The library complexity was confirmed by streaking diluted bacteria onto plates and counting colonies. The total number of colonies was >100x the size of the library which according to simulations in R is far beyond the requirement for faithful replication of a library (data not shown).

Two *Cas9* control viruses were also derived from pKLV2(W-)U6sgRNA5(BbsI)-PGKpuroBFP and pKLV2(gfp)U6sgRNA5(BbsI)-PGKpuroBFP. The new plasmids are correspondingly named pMSCV(W-)U6sgRNA5(BbsI)-PGKpuroBFP and pMSCV(gfp)U6sgRNA5(BbsI)-PGKpuroBFP (Addgene #102797, #102798). The cloning was performed in the same manner as for pMSCV-U6sgRNA(BbsI)-PGKpuro2ABFP.

### Virus production

293T-cells were maintained in Advanced DMEM/F12 (Gibco #12491015) supplemented with geneticin (500ug/ml, Gibco #10131035). At least one day before transfection, cells were kept in media without geneticin. When at roughly 80% confluency (day 1), the cells were transfected using Lipofectamine LTX. To a 10cm dish with 5ml advanced DMEM, we added 3ml OPTI-MEM (Gibco #31985062) containing 36ul LTX, 15ul PLUS (Thermofisher #15338030), and a total of 7.5ug library plasmid and 7.5ug pcl-Eco plasmid58 (Addgene #12371). The OPTI-MEM was incubated for 30 minutes prior to addition. The media was replaced with 5ml fresh Advanced DMEM/F12 the day after transfection (day 2), and virus harvested on day 3. Cells were removed by filtering through a 0.45um syringe filter. Virus was either snap frozen or stored in 4°C (never longer than day 5 before being used).

### Making of mouse strains

Rosa26^Cas9/+^ mice22 were crossed with other mice carrying fluorescent reporters. These strains were Gata3^GFP,59^, Il13^+/Tom,60^ and Il4^tm1.1Wep,61^. For the screens we then pooled mice, both heterozygous and homozygous for *Cas9* expression, male and female, of 8–12 weeks age.

The GATA3-3xFLAG-mCherry mouse strain was produced briefly as follows. The targeting construct was generated by BAC liquid recombineering^62^ such that a CTAP TAG element was linked via a Picornavirus “self-cleaving” T2a peptide^63^ to mCherry red fluorescent protein and placed upstream of a LoxP/Frt flanked promoter driven Neomycin cassette (CTAP-T2a-mCherry-Neomycin). The cassette was flanked by arms of homology and designed to fuse the tagged fluorescent cassette to the terminal Gata3 coding exon, replacing the stop coding and a portion of the endogenous 3’UTR (Supplementary figure 3). Two sgRNAs, left 5’CATGCGTGAGGAGTCTCCAA and right 5’CTTCTACTTGCGTTTTTCGC, were designed to generate double-strand breaks 3’ to the terminal stop codon. The respective complementary oligos (Sigma Genosys) were annealed and cloned into a U6 expression vector.

The targeting construct (2ug), along each U6 guide (1.5*2ug) and wild-type Cas9 (3ug, kind gift from George Church) were nucleofected into 3*10^7^ JM8 F6 C57Bl/6 ES cells using Amaxa Human Stem Cell Kit 2 (Lonza #VPH-5022) and the Amaxa nucleofector B. Subsequent ES cell injections and animal husbandry were carried out by the Sanger Animal facility.

### Validation of *Cas9* mouse

Expression of *Cas9* was confirmed by western blot (anti-Cas9, BioLegend 7A9, #844301) as well as by RT-PCR (primers: cas9_qpcr1/2/r/f). Qualitatively, *Cas9* expression appears to increase during activation of cells (data not included). The function of *Cas9* was also validated using the two control viruses and cytometric analysis. The resulting viruses express both GFP and BFP but only one of them contains a sgRNA targeting its own GFP sequence FACS analysis confirmed a reduction in GFP signal in T-cells infected with the self-targeting virus, as compared to T-cells infected with the control virus (data not included).

### T-cell extraction for CRISPR screening

6-well plates were first prepared at least 2 hours before by adding anti-CD3e (1ul/ml, eBioscences #16-0031-81) in PBS, at least 1.2ml/well, and then kept at 37°C.

Cells were extracted from spleens of up to 30 mice by the following procedure: Spleens were massaged through a 70um strainer into cold IMDM media (strainer slanted to avoid crushing the cells). Cells were spun down at 5min/400g and then resuspended in 5ml red blood cell lysis media (3–4 spleens per 50ml falcon tube). After 4 minutes PBS was added up to 50ml and cells spun again. Cells were then resuspended in cold PBS and taken through a 70um strainer. The cells were counted and spun down again. Finally, the cells were negatively selected using EasySep™ Mouse Naïve CD4+ T Cell Isolation Kit (Stem Cell Technologies, #19765) except for the following modifications: The volume and amount of antibodies were scaled down to 20% of that specified by the manufacturer. Up to the equivalent of the cells of 6–7 spleens can be loaded on one “The Big Easy” EasySep Magnet (Stem Cell Technologies, #18001). Overloading it will cause a severe drop in output cells.

On day 0, the cells were then resuspended in warm IMDM supplemented with 2-Mercaptoethanol “BME” (50 uM Gibco #31350010), IL4 (10ng/ml, R&D Systems 404-ML), IL2 (6ng/ml) and anti-CD28 (3ug/ml, eBioscience #16-0281-86) and Pen/Strep, before being seeded onto the 6-well plates (30–40M cells per plate).

### T-cell culturing for CRISPR screening

On day 1, the cells were infected by the following procedure. To each well, 1.2ml media was added. This media consisted of 80% virus, 20% IMDM, supplemented with BME/IL2/IL4/anti-CD28 at concentrations as before. In addition, the media contained 8ug/ml polybrene. The plate was put in a zip-lock lag and spun at 1100g for about 2 hours at 32°C. The plate was then put in an incubator over-night (never more than 24h in total). The cells in the media were spun down (the cells attached kept in place) and resuspended with media as after the T cell extraction except with the addition of 2ug/ml puromycin. Each well required 3–4ml media. For the 7 day culturing the media had to be replenished after half the day. We estimate that the MOI was about 0.2. The use of puromycin is essential to keep the FACS time down to reasonable levels (commonly 2ng/ml).

### Sorting and genomic DNA extraction

On the day of sorting, cells were extracted and spun down. To eliminate dead cells we performed a “low-g spin”, 5 minutes at 200g. This brought the viability up to roughly 50%. We have in addition tried other methods such as Ficoll (works slightly better but takes 30 minutes and is harder to reproduce) and Miltenyii Dead Cell Removal Kit. In our experience, the Miltenyii kit works great on uninfected cells but effectively removed almost every infected cell when attempted on the real sample. This is most likely because the kit does negative selection against Annexins which might be promoted by the virus or the puromycin.

In the cases when we used antibody reporters, we first fixed and permeabilized using the Foxp3 Transcription Factor Staining Buffer Kit (eBioscience, #00-5523-00). We then used the following antibodies: PE Mouse anti-XBP-1S (BD Biosciences, #562642), FITC anti-IRF4 (BD Biosciences, #11-9858-80) and Alexa Fluor 488 Mouse anti-GATA3 (BD Biosciences, #560077).

For sorting, cells were resuspended at 40M/ml in IMDM with BME and 3mM EDTA (PBS for the stained cells). The use of EDTA is essential to ensure singlet events at this high cell concentrations. The cells were then sorted into IMDM using either a Beckman MoFlo or MoFlo XDP, or BD Influx. For non-stained screens we could use BFP to ensure that the cells passed were infected. For the stained screens the BFP signal was disrupted by the staining and we performed it blindly. The subsequent steps are not affected by the addition of uninfected cells. During protocol development, the FACS data was analysed using the software FACSanadu (http://www.facsanadu.org).

After sorting the cells, we performed DNA extraction in two different ways. When using fluorescent reporter strains we used the Blood & Cell Culture DNA Midi Kit (Qiagen #13343).

For the fixed cells, due to lack of suitable commercial kits (The FFPE kits we have seen are for low amounts of DNA only), we instead performed DNA extraction as follows. Sorted cells were pellet using a table-top centrifuge at 2000g, 5 minutes. Cell pellet was resuspended in 500 ul Lysis Buffer I (50 mM HEPES.KOH, pH 7.5, 140 mM NaCl, 1 mM EDTA, 10% Glycerol, 0.5% NP-40, 0.25% Triton X-100) and rotate at 4°C for 10 minutes. Cells were spun down at 2000g, 5 minutes, resuspended in 500 ul Lysis Buffer II (10 mM Tris.Cl, pH 8.0, 200 mM NaCl, 1 mM EDTA, 0.5 mM EGTA) and rotate at 4°C for 10 minutes. Then the cells were pelleted again at 2000g for 5 minutes, and the cell pellet was resuspended in 25 ul Lysis Buffer III (Tris.Cl, pH 8.0, 100 mM NaCl, 1 mM EDTA, 0.5 mM EGTA, 0.1% Na-Deoxycholate, 0.5% N-lauroylsarcosine). Then 75 ul TES Buffer (50 mM Tris.Cl pH 8.0, 10 mM EDTA, 1% SDS) was added to the cell suspension. This 100 ul reaction was put on a thermomixer to reverse crosslinking at 65°C, overnight. Then 1 ul proteinase K (20 mg/ml, ThermoFisher #100005393) was added, and protein was digested at 55°C for 1 hour. DNA was purified using MinElute PCR purification kit (Qiagen) according to the manufacturer’s instruction. DNA concentration was measured by a Nanodrop.

### Sequencing of CRISPR virus insert

The genomic DNA was first PCR-amplified (primers: gLibrary-HiSeq_50bp-SE_u1/l122) in a reaction with Q5 Hot Start High-Fidelity 2X Master Mix (NEB #M0494L). In each 50ul reaction, we loaded up to 3ug DNA. From each reaction we pipetted and pooled 5ul, before purifying it using the QIAquick PCR purification kit (Qiagen #28104). The purified product was then further PCR-amplified using KAPA HiFi HotStart ReadyMix (Kapa #KK2602) and iPCRtag sequencing adapters64. After Ampure XP bead purification (beads made up 70% of the solution) and Bioanalyzer QC, the libraries were pooled and sequenced with a HiSeq 2500 (Illumina #SY-401–2501, 19bp SE). The custom primers U6-Illumina-seq2 (R1) and iPCRtagseq (index sequencing) were used for this purpose. The original sgRNA library contained 86,035 distinct sgRNAs. In a representative sequencing run (*Gata3*, using antibody selection) the sgRNAs with fewer than 500 reads encompassing 91% of the total complexity.

### Analysis and QC of CRISPR hits

Sequencing BAM-files were transformed into FASTQ using samtools and bamToFastq. A custom Java program was then used extract per-sgRNA read counts. From these, per-gene p-values were calculated using MAGeCK^28^ using the positive and negative cell fraction from each screen. The hit rankings were then compared using R. To obtain a total per-gene score, we first calculate the total rank from one screen as r=min(r_pos_,r_neg_), using the ranks from the positive and negative enrichments respectively. Then, to calculate the composite score of two or more screens, we used the geometric mean (r_1_r_2_r_3_…r_n_)^1/n^. Follow-up hits were manually picked as those scoring high between the replicates, with genes of low expression level qualitatively filtered out using ImmGen^65^.

The BaIOPSE model was implemented in STAN^66^ using the RStan interface. For the full model implementation and parameters, with variances rather defined by the exponentials over the priors, we refer to the source code. 12 Markov chains were run 800 steps and convergence was checked by the r-value. The top 300 hits were used to calculate GO term *p*-values. GO terms were obtained in R by GO.db^67^ and assessed individually using a Fisher exact test.

### Mouse time-course RNA-seq

CD4^+^CD62L^+^ naive T cells were purified from spleens of wild-type C57BL/6JAX adult (6 – 8 weeks) mice using the CD4+CD62L+ T Cell Isolation Kit II (Miltenyii #130-093-227). Cell culture plates were coated with anti-CD3e antibody (1 ug/ml, eBioscences #16-0031-81) in 1X DPBS (Gibco) at 4°C overnight. Purified naive T cells were seeded at a concentration of 1 M cells/ml on the coated plates in IMDM (Gibco) with 10% heat-inactivated FBS (Sigma #F9665-500ML), supplied with 5 ug/ml anti-CD28 (eBioscience #16-0281-86) with (Th2) or without (Th0) 10 ng/ml mouse recombinant IL-4 (R&D Systems #404-ML-050). Cells were cultured in plates for up to 72 hours.

Total RNA was purified by Qiagen RNeasy Mini Kit according to manufacturer’s instruction, and concentration was determined by a Nanodrop. A total of 500 ng RNA was used to prepare sequencing libraries using KAPA Stranded mRNA-seq Kit (KAPA #07962193001) according to manufacturer’s instructions. Sequencing was performed on an Illumina HiSeq 2000 (125bp PE, v4 chemistry).

### Human time-course RNA-seq

Mononuclear cells were isolated from the cord blood of healthy neonates at Turku University Central Hospital using Ficoll-Paque PLUS (GE Healthcare, #17-1440-02). CD4+ T cells were then isolated using the Dynal bead-based positive isolation kit (Invitrogen). CD4+ cells from three individual donors were activated directly in 24w plates with plate-bound anti-CD3 (500ng/well, Immunotech) and soluble anti-CD28 (500 ng/mL, Immunotech) at a density of 2×10^6^ cells/mL of Yssel’s medium68 containing 1% human AB serum (PAA). Th2 cell polarization was initiated with IL-4 (10 ng/ml, R&D Systems). Cells activated without IL-4 were also cultured (Th0).

### Time-course ATAC-seq data generation

Experiments were done according to the published protocol^69^ with some modification. Briefly, 50,000 cells were washed with ice cold 1X DPBS twice, and resuspended in a sucrose swelling buffer (0.32 M sucrose, 10 mM Tris.Cl, pH 7.5, 3 mM CaCh, 2 mM MgCb, 10% glycerol). The cell suspension was left on ice for 10 minutes. Then, a final concentration of 0.5% NP-40 was added, and the cells suspension was vortexed for 10 seconds and left on ice for 10 minutes. Nuclei was pelleted at 500 g at 4°C for 10 minutes. Nuclei were washed once with 1X TD buffer (from Nextera DNA Library Preparation Kit, Illumina, #FC-121-1030), and resuspended in 50 ul tagmentation mix containing:

- 25 ul 2X TD buffer (Nextera DNA Library Preparation Kit, Illumina #FC-121-1030)
- 22.5 ul H_2_O
- 2.5 ul TDE1 (Nextera DNA Library Preparation Kit, Illumina #FC-121-1030)

The tagmentation reaction was carried out on a thermomixer at 37°C, 800 rpm, for 30 minutes. The reaction was stopped by the addition of 250 ul (5 volumes) Buffer PB (from Qiagen MinElute PCR Purification Kit), The tagmented DNA was purified by Qiagen PCR Purification Kit according to manufacturer’s instructions and eluted in 12.5 ul Buffer EB from the kit, which yielded ~10 ul purified DNA.

The library amplification was done in a 25 ul reaction include:

- 10 ul purified DNA (from above)
- 2.5 ul PCR Primer Cocktail (Nextera DNA Library Preparation Kit, Illumina #FC-121-1030)
- 2.5 ul N5xx (Nextera index kit, Illumina #FC-121-1012)
- 2.5 ul N7xx (Nextera index kit, Illumina #FC-121-1012)
- 7.5 ul NPM PCR master mix (Nextera DNA Library Preparation Kit, Illumina #FC-121-1030)

PCR was performed as follows:

- 72°C 5 minutes
- 98°C 2 minutes
- [98°C 10 secs, 63°C 30 secs, 72°C 60 secs] × 12
- 10°C hold

Amplified libraries were purified by double Agencourt AMpureXP beads purifications (Beckman Coulter, #A63882). 0.4X beads:DNA ratio for the first time, flow through was kept (removing large fragments); 1.4X beads:DNA ratio for the second time, beads were kept. Libraries were eluted from the beads by elution in 20 ul Buffer EB (from Qiagen PCR Purification Kit).

1 ul library was run on a Agilent Bioanalyzer to check size distribution and quality of the libraries.

Sequencing was done with an Illumina Hiseq 2500 (75 bp PE).

### ChIP-seq data generation

ChIPmentation^70^ was used to investigate the TF binding sites. 1 million cells from each sample were crosslinked in 1% HCHO (prepared in 1X DPBS) at room temperature for 10 minutes, and HCHO was quenched by the addition of glycine at a final concentration of 0.125 M. Cells were pelleted at 4°C at 2000 x g, washed with ice-cold 1X DPBS twice, and snapped frozen in liquid nitrogen. The cell pellets were stored in−80°C until the experiments were performed. ChIPmentation was performed according to the version 1.0 of the published protocol (http://www.medical-epigenomics.org/papers/schmidl2015/) with some modifications at the ChIP stage. The antibody used were IRF4 (sc-6059), BATF (sc-100974) and FLAG (Sigma M2, #F3165).

Briefly, cell pellets were thawed on ice, and lysed in 300 ul ChIP Lysis Buffer I (50 mM HEPES.KOH, pH 7.5, 140 mM NaCl, 1 mM EDTA, pH 8.0, 10% Glycerol, 0.5% NP-40, 0.25% Triton X-100) on ice for 10 minutes. Then cells were pelleted at 4°C at 2000 x g for 5 minutes, and washed by 300 ul ChIP Lysis Buffer II (10 mM Tris.Cl, pH 8.0, 200 mM NaCl, 1 mM EDTA, pH 8.0, 0.5 mM EGTA, pH 8.0), and pelleted again at 4°C at 2000 x g for 5 minutes. Nuclei were resuspended in 300 ul ChIP Lysis Buffer III (10 mM Tris.Cl, pH 8.0, 100 mM NaCl, 1 mM EDTA, 0.5 mM EGTA, 0.1% Sodium Deoxycholate, 0.5% N-Lauroylsarcosine). Chromatin was sonicated using Bioruptor Pico (Diagenode) with 30 seconds ON/30 seconds OFF for 10 cycles. 30 ul 10% Triton X-100 was added into each sonicated chromatin, and insoluble chromatin was pelleted at 16,100 x g at 4°C for 10 minutes. 1 ul supernatant was taken as input control. The rest of the supernatant was incubated with 10 ul Protein A Dynabeads (Invitrogen) pre-bound with 1 ug anti-FLAG in a rotating platform in a cold room overnight. Each immunoprecipitation (IP) was washed with 500 ul RIPA Buffer (50 mM HEPES.KOH, pH 7.5, 500 mM LiCl, 1 mM EDTA, 1% NP-40, 0.7% Sodium Deoxycholate, check components) for 3 times. Then, each IP was washed with 500 ul 10 mM Tris, pH 8.0 twice, and resuspended in 30 ul tagmentation reaction mix (10 mM Tris.Cl, pH 8.0, 5 mM Mg2Cl, 1 ul TDE1 (Nextera)). Then, the tagmentation reaction was put on a thermomixer at 37°C for 10 minutes at 800 rpm shaking. After the tagmentation reaction, each IP was washed sequentially with 500 ul RIPA Buffer twice, and 1X TE NaCl (10 mM Tris.Cl, pH 8.0, 1 mM EDTA, pH 8.0, 50 mM NaCl) once. Elution and reverse-crosslinking were done by resuspending the beads with 100 ul ChIP Elution Buffer (50 mM Tris.Cl, pH 8.0, 10 mM EDTA, pH 8.0, 1% SDS) on a thermomixer at 65°C overnight, 1,400 rpm. DNA was purified by MinElute PCR Purification Kit (QIAGEN, #28004 and eluted in 12.5 ul Buffer EB (QIAGEN kit, #28004), which yielded ~10 ul ChIPed DNA.

The library preparation reactions contained the following:

- 10 ul purified DNA (from above)
- 2.5 ul PCR Primer Cocktails (Nextera DNA Library Preparation Kit, Illumina #FC-121-1030)
- 2.5 ul N5xx (Nextera Index Kit, Illumina #FC-121-1012)
- 2.5 ul N7xx (Nextera index kit, Illumina #FC-121-1012)
- 7.5 ul NPM PCR Master Mix (Nextera DNA Library Preparation Kit, Illumina #FC-121-1030)

PCR was set up as follows:

- 72°C, 5 mins
- 98°C, 2 mins
- [98°C, 10 secs, 63°C, 30 secs, 72°C, 20 secs] × 12
- 10°C hold

The amplified libraries were purified by double Ampure×P beads purification: first with 0.5× bead ratio, keep supernatant, second with 1.4× bead ratio, keep bound DNA. Elution was done in 20 ul Buffer EB (QIAGEN).

1 ul of library was run on an Agilent Bioanalyzer to see the size distribution. Sequencing was done on an Illumina Hiseq 2000 platform (75 bp PE, v4 chemistry).

### ChIP-seq peak analysis

The reads were first trimmed using Trimmomatic 0.36^71^ with settings ILLUMINACLIP:NexteraPE-PE.fa:2:30:10 LEADING:3 TRAILING:3 SLIDINGWINDOW:4:15 MINLEN:30. Peaks were then called using MACS2^44^, merged over time, and annotated using HOMER^72^.

The quality of the peaks was assessed using the two available replicates for each time point. While the trend over time agreed, the number in each time point did not. For this reason we decided to consider the union of the peaks rather than the common peaks.

The sequences of the detected ChIP-seq peaks were extracted using “bedtools getfasta”^73^, for 200, 300, 400, 500bp regions around the peaks. These were fed into MEME^38^ for additional motif discovery.

### Time-course RNA-seq differential expression

Gene expression from RNA-seq data was quantified in TPM using Salmon v0.6.0^74^, with the parameters --fldMax 150000000 --fldMean 350 --fldSD 250 --numBootstraps 100 --biasCorrect --allowOrphans --useVBOpt. The cDNA sequences supplied contain genes from GRCm38 (mouse), GRCh38 (human) and sequences from RepBase, as well as ERCC sequences and an eGFP sequence.

Differentially expressed (DE) genes were found using the Sleuth R package^75^, using the wasabi R package (https://github.com/COMBINE-lab/wasabi) to allow it to accept Salmon input data. To strengthen the test of differential dynamics between Th2 and Th0 culture conditions, instead of testing each time point individually (with few replicates), we separated time into early (<=6h) and late (>6h). The DE test consisted of a likelihood-ratio test using the sleuth_lrt function, where the full model contained terms accounting for the culture condition, for the temporal effect (modelled as a spline with 5 degrees of freedom) and for an interaction of both terms. To capture the Th0/Th2 difference, the reduced model only contained a term accounting for the time variation, modelled as before. A gene is considered differentially expressed for p-value < 0.01

### Human/Mouse *Stat6* comparison

Targets of Stat6 and Il4 as defined by time-course microarray and ChIP-seq data was downloaded from a previous study^5^.

### ATAC-seq motif extraction

ATAC-seq reads were aligned using Bowtie 2^76^ with the parameter −X 2000 and the mouse genome mm10. This was followed by peak calling on each replicate individually using MACS2^44^ with the function “callpeak” and the parameters −B --SPMR --call-summits. The peaks obtained were kept if they overlapped a peak from the other replicate of the same time point by at least 50%. In these cases, the new peak would equal the combined coordinates of all the overlapping peaks considering all replicates and time points.

Peaks were classified (annotatePeaks.pl --annStats) as intronic, exonic, upstream or intergenic, according to the gene feature they intersected. Intersection is scored first considering the number of bases overlapped, and then the closeness in size between the peak and the feature.

Known motif detection was performed on the peaks’ sequences using FIMO^45^, and motifs from the JASPAR 2016 database^77^ considering only those starting in MA or PB. In addition, we supplemented with a more recent list of C2H2 motifs^78^. To make the analysis more targeted, only motifs from TFs DE between Th2 and Th0 were considered, and for each of them a single motif was selected, prioritizing the longest ones with the lowest mean entropy.

The overlap between human and mouse was calculated using liftOver-minMatch=0.03-multiple. Roughly 100 peaks mapped to multiple sites and were thus ignored. LiftOver was also performed on individual TF sites from FIMO. The overlap between organisms was calculated using R GenomicRanges^79^. The overlap procedure was done at the peak and detected motif levels.

We found that the analyses throughout the paper appear to give similar results when using all mouse peaks as opposed to only using the conserved (overlapping) peaks. However, the ChIP-seq peaks of GATA3, IRF4 and BATF appear more comparable to ATAC-seq predicted sites if only the conserved sites are used are used in the MARA.

### ATAC-seq chromatin dynamics analysis

The height of the peaks, as well as any reads outside the peaks, were quantified using bedtools73. The peak levels were divided by the background signal for normalization. Further, to make the contributions from different peaks comparable, they were normalized to the level of the second time point. The contribution of motifs over time is defined as the average peak signal in which they are present.

### UCSC visualization of ChIP-seq and ATAC-seq

The MACS2 BedGraph files were prepared for UCSC visualization using bedSort and bedGraphToBigWig.

### ARACNe analysis

The ARACNe-AP^51^ package was used as follows. For a given network, the RNA-seq data is stored in files net.exp. The list of core genes (TFs and cytokines) is stored in net.tf. To initialize the algorithm, this command is used: “java-Xmx5G-jar Aracne.jar-e net.exp-o net.out --tfs net.tf --pvalue 1E-5 --seed 1 --calculateThreshold --nodpi”. Next, the mutual information (MI) calculation is done using “java-Xmx7G-jar Aracne.jar-e net.exp-o net.out --tfs net.tf --pvalue 1E-5 --seed…. --nodpi”. Finally, the data is summarized using “java-Xmx120000M-jar Aracne.jar-o net.out --tfs net.genes --consolidate --nodpi”.

Analysis of the network was performed in R. To validate the network the clustering of hits in a certain screen Z was assessed based on the number of neighbours X, Y, both being Z-hits. Using a bootstrap test, the number of such neighbours was compared to randomized networks where the identity of the vertices was swapped.

To check if a gene X is especially important for Z, enrichment for nearby Z-hits was tested. For each gene X (as a vertex), the other genes (vertices) one edge apart were checked for being Z-hits. The gene X is considered enriched for hits if it has more hits than average, tested using a binomial test.

### MARA analysis

The MARA analysis was performed as follows. Early and late times were analyzed independently. For each of the two durations, the connectivity matrix was constructed based on if a motif peak was present for a gene at any time. The number of such peaks, ignoring time fluctuations, were entered as the connectivity value. The full RNA-seq time-course data for either Th0 or Th2 was used as the signal. These two files were uploaded to ISMARA53 using expert mode.

In the MARA comparison over time, Th0 and Th2 difference is calculated as the average MARA activity difference over time. The activity increase is taken as the difference in activity at the first and last time points for Th0.

### Follow-up knock-out RNA-seq data generation

The backbone pMSCV-U6sgRNA(BbsI)-PGKpuro2ABFP was digested using BbsI and purified on a gel. 96*2 desalted primers for the sgRNA insert were obtained from Sigma in premixed and diluted format. They were diluted to 10uM in T4 ligation buffer (NEB, #M0202T) and annealed (cooling from 98°C to 4°C during 1 hour on a PCR block). Ligations were performed in 10ul volume, in a 96w PCR on ice. Transformed *E. coli* (Stbl3, made competent in lab) were streaked onto 10cm ampicillin agar plates using an 8 channel pipette.

To avoid validating individual colonies, a mixture of at minimum 10+ colonies were picked and mixed for each clone. Digest by BbsI of a few representative shows at the minimum presence of clones without original bbsI spacer. Bacteria were grown overnight in a 96w deep-well plate having an air-permeable seal. Minipreps were made using a homemade gravity manifold holding miniprep tubes (blueprint for laser cutting available on request). The virus was subsequently made in 293T-cells, in 24w format. The virus was then harvested into a 96w deep-well plate and any 293T removed by centrifugation.

Naive T cells were extracted from 3 mice independently, this time with the Naive CD4+ T Cell Isolation Kit (Miltenyii #130-104-453) according to manufacturer’s instruction. Cells were seeded at 200k/well density in 96w format. Infection and puromycin selection was then performed as before. On day 5, cells were washed and dead cells removed by low-G spin. This typically raised the viability from roughly 10–20% to 60% according to Trypan blue. Cells were spun down and as much of the media removed as possible. Up to 100ul of buffer RLT+ was then added to each well and plates frozen. Later, plates were thawed and RNA extracted by adding 100ul of Ampure XP beads. Purification was done by a robot, with 2x200ul EtOH wash and final suspension in RNAse-free water. RNA was then diluted to 500ng/ul and 2ul was taken as input into non-capping DogSeq (manuscript in preparation). This protocol is for this application roughly equivalent to Smartseq-2^80^. Libraries were made using Nextera XT and all 96*3 libraries sequenced with a HiSeq 2500 (150bp PE).

### Follow-up knock-out RNA-seq analysis

Reads were filtered using Cutadapt for the Smart-seq2 TSO and mapped using GSNAP^81^. The software featureCounts was then used to produce a final count table^82^. The effects of the KO was studied using an EdgeR^83^ linear regression model using the KO with scrambled sgRNA as reference point. We studied the impact of the virus infection level, measured as a function of BFP, and found it to be confounding. To obtain stronger DE effect for future KO experiments we recommend that non-infected cells are removed by FACS sorting rather than puromycin selection. Individual replicates were compared in terms of p-value and correlation of DE genes when one sgRNA was used *versus* when several sgRNAs targeting the same gene were pooled. Libraries with low replicability or low virus infection were manually removed.

We define a differentiation axis as the DE genes (using DEseq2) from the RNA-seq time-course, Th0 vs Th2 at t=72h, with p<10^−10^. The activation axis is similarly defined as the DE genes Th2 at t=0h vs Th2 at t=72h, with p<10^−10^.

Similarly, we define axes for each T helper type using DE data from Th-express^46^ with p<10^−2^. For Figure 6ab we then calculate the similarity score as s_i_=Σ_i∈g_ ref_i_, ko_i_/|g|, where g is the set of genes which in the KO have a 2-fold change.

The plasticity dimension reduction Figure 5c is produced as follows. A standard pinwheel type diagram cannot be produced because of a lack of absolute T cell type references (cultured under the same conditions as the KO, with scrambled sgRNA, and same RNA-seq protocol). The diagram was instead based on the previous Th-Express DE vectors. The position of each KO is calculated as αΣS_i_**v**_i_ where **v** is the vector from Th2 to Thi, and α an arbitrary constant. As a result, the labels Thi are merely directions toward T helper cell types, and not absolute coordinates in the cell type state space.

